# Novel cell states arise in embryonic cells devoid of key reprogramming factors

**DOI:** 10.1101/2024.05.13.593729

**Authors:** Scott E. Youlten, Liyun Miao, Caroline Hoppe, Curtis W. Boswell, Damir Musaev, Mario Abdelmessih, Smita Krishnaswamy, Valerie A. Tornini, Antonio J. Giraldez

## Abstract

The capacity for embryonic cells to differentiate relies on a large-scale reprogramming of the oocyte and sperm nucleus into a transient totipotent state. In zebrafish, this reprogramming step is achieved by the pioneer factors Nanog, Pou5f3, and Sox19b (NPS). Yet, it remains unclear whether cells lacking this reprogramming step are directed towards wild type states or towards novel developmental canals in the Waddington landscape of embryonic development. Here we investigate the developmental fate of embryonic cells mutant for NPS by analyzing their single-cell gene expression profiles. We find that cells lacking the first developmental reprogramming steps can acquire distinct cell states. These states are manifested by gene expression modules that result from a failure of nuclear reprogramming, the persistence of the maternal program, and the activation of somatic compensatory programs. As a result, most mutant cells follow new developmental canals and acquire new mixed cell states in development. In contrast, a group of mutant cells acquire primordial germ cell-like states, suggesting that NPS-dependent reprogramming is dispensable for these cell states. Together, these results demonstrate that developmental reprogramming after fertilization is required to differentiate most canonical developmental programs, and loss of the transient totipotent state canalizes embryonic cells into new developmental states *in vivo*.

## Introduction

After fertilization, the embryo undergoes a large-scale reprogramming of the oocyte and sperm nucleus and the cytoplasm into a transient totipotent state. Following this initial reprogramming during the activation of the zygotic program and the clearance of the maternal mRNAs, embryonic cells progressively lose this transient totipotent state as they differentiate into terminal, differentiated cell states.

In 1957, Waddington used a model of a ball rolling down a landscape through a series of canals to describe the progressive restriction and determination of cell fate during development (Waddington, 1957). As time goes on, the landscape goes through a succession of bifurcations, giving rise to new valleys and new possible cell fates (Ferrell, 2012). In this model, intrinsic and extrinsic factors govern a cell’s state and its probabilistic journey down progressive slopes and hills, with each previous cell state priming a cell towards a “canal” or path iteratively until it reaches its final state (Waddington, 1957; Ferrell, 2012). The following year, this model was challenged by somatic cell nuclear transfer, wherein a somatic nucleus transplanted into an oocyte can be reprogrammed to a totipotent state by factors in the cytoplasm (Gurdon, Elsdale and Fischberg, 1958). Decades later it was demonstrated that reprogramming could be achieved by reintroducing cocktails of transcription factors (TF) involved in epigenetic reprogramming during development, including the homeobox TF Nanog, the POU Class 5 Homeobox TF Pou5f1 (known previously as Oct4) and the SRY-Box TF Sox2 (Takahashi and Yamanaka, 2006; Yu *et al*., 2007). Yet, the efficiency with which induced pluripotent cells give rise to naturally occurring somatic cell fates is orders of magnitude below that of embryonic stem cells, and it is unclear if this is an intrinsic deficit in reprogramming itself or if it is due to the absence of extrinsic environmental cues. Recent single cell analyses comparing mutant or perturbed states to wild type (WT) states has found that cell fates remained restricted to known cell states in WT embryos (Farrell *et al*., 2018; Wagner *et al*., 2018; Saunders *et al*., 2023). In the context of Waddington’s landscape, these findings suggest that mutant cells do not diverge to create *de novo* gene expression programs and new differentiation paths; instead, their states converge or canalize into existing states found in developing wild type cells.

In zebrafish, Nanog, Pou5f3, and Sox19b (NPS) (functionally analogous to mammalian Nanog, Pou5f1 and Sox2, respectively) are required during the maternal-to-zygotic transition (MZT) for the first nuclear reprogramming event and subsequent zygotic genome activation (ZGA) (Chambers and Tomlinson, 2009; Onichtchouk, 2012; Lee *et al*., 2013; Sukparangsi *et al*., 2022). These pioneer factors reprogram the nucleus by selectively opening chromatin and coordinating the first waves of zygotic gene expression. They also reprogram the cytoplasm by activating the conserved microRNA *miR*-*430,* which represses and degrades many maternally deposited transcripts to complete the transition to the zygotic program (Giraldez *et al*., 2006; Yartseva and Giraldez, 2015).

We recently generated a triple maternal-zygotic *nanog*-/-;*pou5f3*-/-;*sox19b*-/- mutant zebrafish (MZ*nps*) devoid of all three pioneer factors (Miao *et al*., 2022). MZ*nps* embryos display defects in reprogramming compared to WT, including dramatic deficiencies in chromatin accessibility, failure to activate a fraction of the zygotic genome, including *miR-430* for maternal program clearance, resulting in developmental arrest before gastrulation at 4 hours post fertilization (hpf) (Lee *et al*., 2013; Miao *et al*., 2022). Given that the MZ*nps* embryo mutant environment cannot sustain their development, it is unclear whether their failure to reprogram precludes these embryonic cells from differentiating into any cell types. These mutants now allow us to test the nuclear and cytoplasmic reprogramming hypothesis, and whether new cellular states or developmental canals can be introduced in the Waddington landscape of embryogenesis if the process of nuclear and cellular reprogramming are affected.

In this study, we ask a fundamental question in developmental biology: what is the intrinsic differentiation potential of embryonic cells that lack the first critical step of developmental reprogramming into a transient totipotent state? By transplanting MZ*nps* cells into a WT environment, we find evidence that cells that fail to reprogram can diverge to create new canals by acquiring distinct, novel cell states in development.

## Results

### Investigating the role of NPS in cell differentiation using single-cell RNA sequencing

Embryos lacking MZ*nps* arrest at 4 hpf and fail to gastrulate, precluding the analysis of their function in cell differentiation beyond genome activation (Lee *et al*., 2013; Miao *et al*., 2022). To understand whether MZ*nps* mutant cells differentiate into different cell types, we transplanted GFP+ labeled cells from either mutant MZ*nps* or control WT donor embryos into WT host embryos at the blastula stage (∼3.3 hpf) (illustrated in Figure 1A) and imaged host embryos cells at early organogenesis (∼12 hpf) (Figures 1B-1D). We found both MZ*nps* and WT labeled donor cells in WT hosts at 12 hpf, albeit with less morphological differentiation of MZ*nps* cells compared to WT donor cells (Figure 1B-D), indicating that a subset of MZ*nps* mutant cells integrate in the WT host during organogenesis without the first reprogramming step.

**Figure 1.**
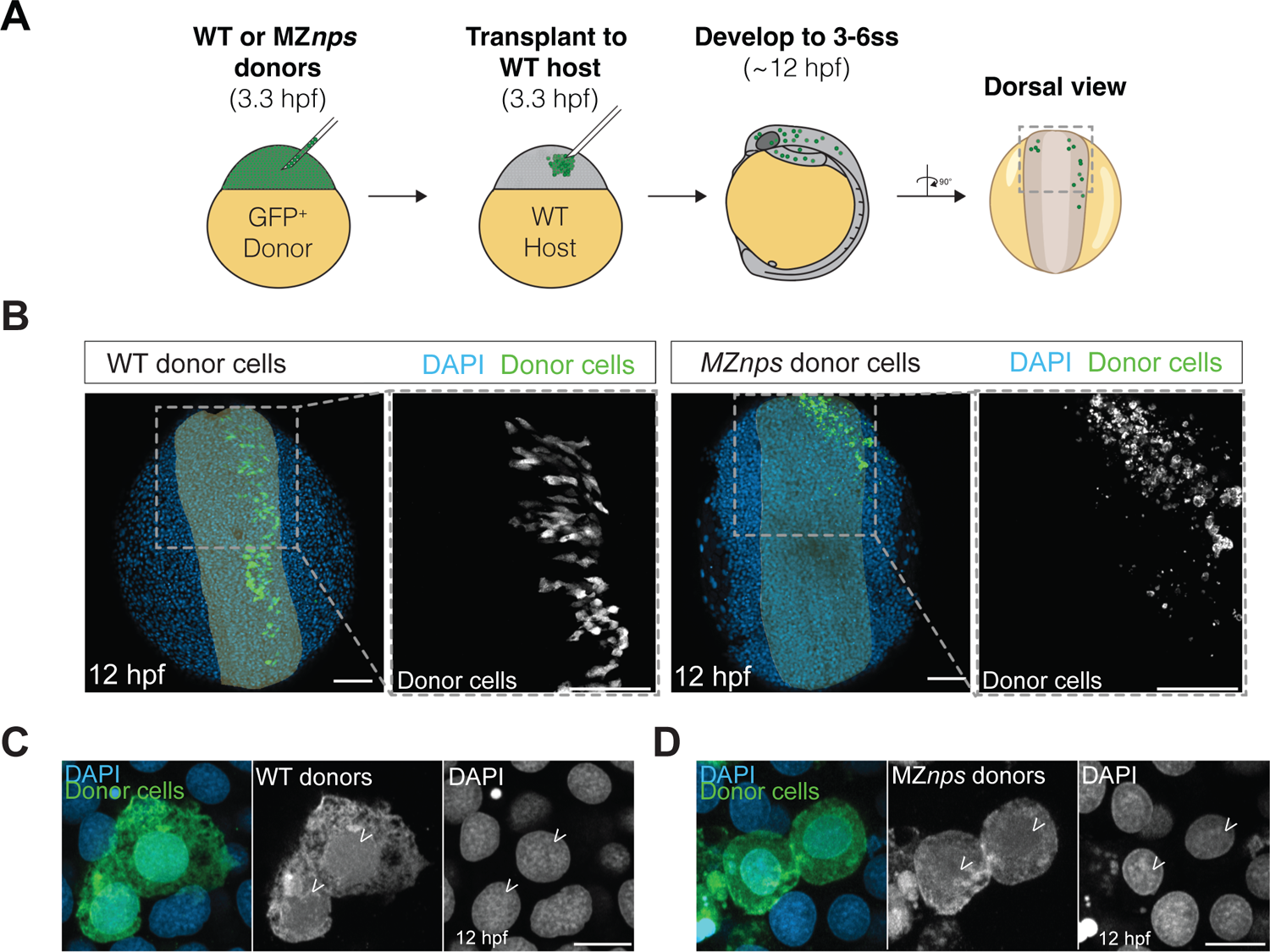
Embryonic cells lacking pioneer factors Nanog, Pou5f3, and Sox19b can survive in a wild-type embryo A. Schematic illustration of zebrafish blastula transplants. Wild type (WT) or MZ*nps* donor embryos were injected at the one-cell stage with GFP mRNA. At 3.3 hours post-fertilization (hpf), GFP-labeled donor cells from either WT or MZ*nps* embryos were transplanted into unlabeled WT hosts imaged at the 3-6 somite stage (∼12 hpf). B. WT host embryos imaged at 3-6SS containing GFP+ donor cells (green) from either WT (left) or *MZnps* mutant embryos (right) that were transplanted at the 1-cell stage. Embryos are oriented dorsally with the region of analysis indicated in grey. Nuclei are labelled with DAPI (blue). Insets show GFP+ cells (gray). Scale bars = 100 mm. C. Representative example of GFP+ WT donor cells (green) showing “normal morphology”. Nuclei of WT host and donor cells are labelled with DAPI (blue). Scale bar, 10 μm. D. Representative example of GFP+ MZ*nps* donor cells (green) showing “normal morphology”. Nuclei of WT host and donor cells are labelled with DAPI (blue). Scale bar, 10 μm.

To investigate the cellular identity of the MZ*nps* cells, we developed a method for rapid enrichment of donor cells relative to host cells and subsequent single-cell RNA sequencing (scRNA-seq) (illustrated in Figure 2A) (Miltenyi *et al*., 1990; Wattrus and Zon, 2020). To this end, we expressed a truncated human cell-surface marker CD4 (hCD4) and DsRed in donor WT or MZ*nps* embryos using mRNA injection. Briefly, we transplanted labeled WT or MZ*nps* donor cells into WT host embryos at 3.3 hpf, allowed them to develop, and dissociated embryos at 3-6 somite stage (∼12 hpf). We enriched the dissociated cell suspensions 2.26-fold with hCD4-binding MicroBeads and magnetic column retention (Figure S1A-1D). We then performed single-cell RNA sequencing (10x Genomics) on donor-enriched cell suspensions.

**Figure 2.**
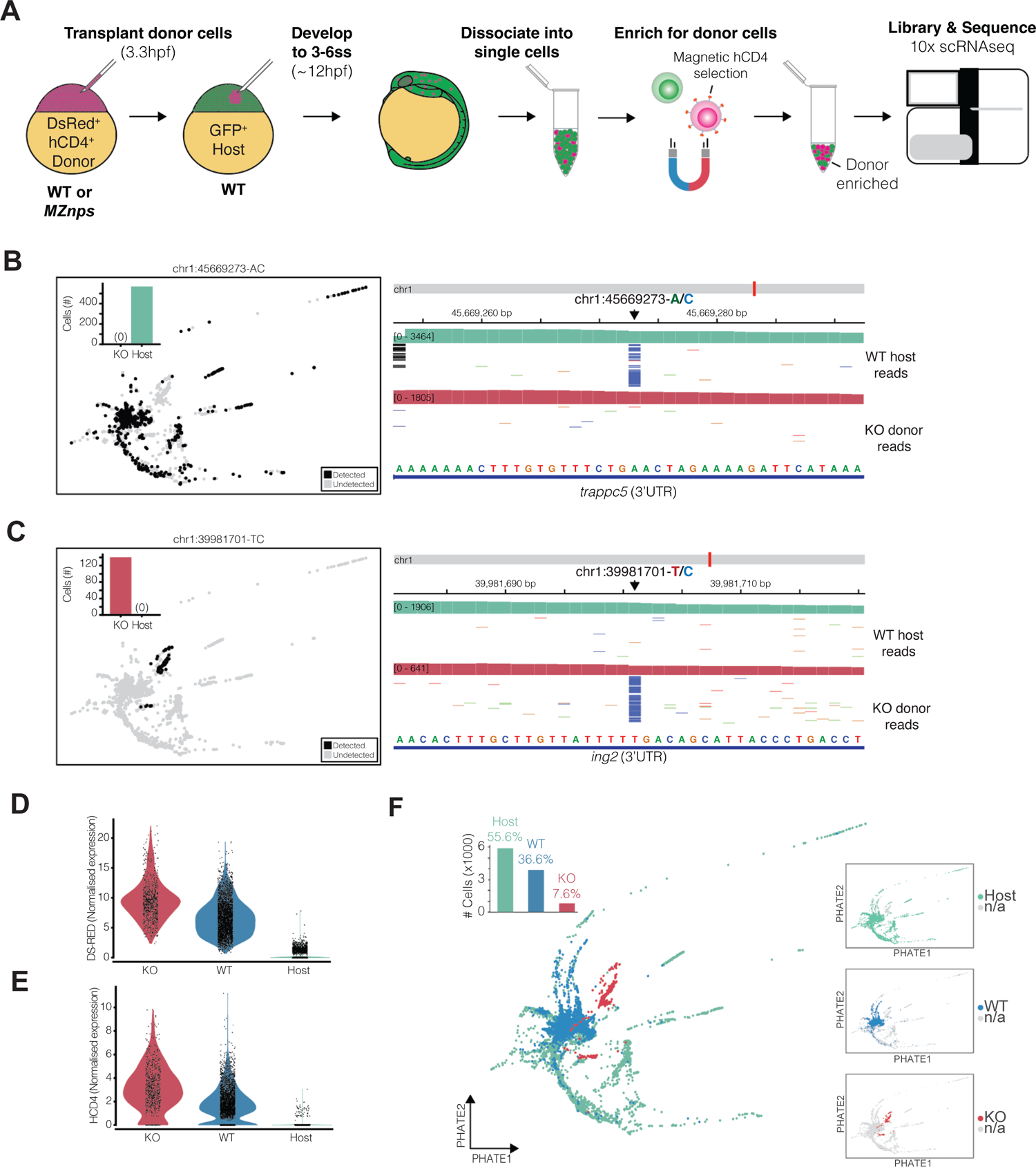
Enrichment and identification of *MZnps* donor cells enables *in vivo* profiling of cell states **A.** Schematic of approach used to enrich cell-suspensions for donor cells from dissociated host embryos. mRNA encoding the truncated human cell-membrane protein CD4 (hCD4) was injected into donor zebrafish embryos at the once-cell stage. Cell suspensions were then incubated with magnetic anti-CD4 MicroBeads allowing enrichment of hCD4+ cells. Single-cell sequencing was performed on the donor-cell enriched suspension. **B.** The detection per cell (left) and read-data alignment (right) for an A to C mutation in the 3’UTR of *trappc5* (chr1:45669273), present in reads from WT host cells (top) and not MZ*nps* donor cells (bottom). Bar plots show the number of cells in which the SNV was detected per condition. **C.** The detection per cell (left) and read-data alignment (right) for an T to C mutation in the 3’UTR of *ing2* (chr1: 39981701), only detected in reads from MZ*nps* donor cells (bottom). Bar plots show the number of cells in which the SNV was detected per condition. **D.** Violin plot of normalized DS-Red mRNA expression in single cells from *MZnps* (KO) donor cells (red) or WT donor cells (blue) and WT host cells (green). **E.** Violin plot of normalized hCD4 mRNA expression in single cells from *MZnps* (KO) donor cells (red) or WT donor cells (blue) and WT host cells (green). **F.** A 2D PHATE embedding of WT host (green), WT donor (blue), and MZ*nps* (red) cells overlaid and individually (inset). Barplot shows the number and relative percentage of cells per condition in the total datasets.

We analyzed gene expression in 10,551 cells across 19,803 genes after quality control (see Methods). Among those, we identified 810 MZ*nps* mutant donor cells across two independent replicates and 3869 WT donor cells based on the enrichment of DsRed and hCD4 marker-gene sequences and the expression of strain specific single nucleotide variants (SNV) in each population (Figures 2B-2E, S1E and S1F). The remaining cells belonged to the WT host (Figure 2F). Transcriptome analysis of MZ*nps* and WT cells revealed that the majority of zygotic genes (17775) were eventually activated in both populations with similar numbers of genes detected (Figure S1G), including 89.9% of zygotic genes common to both populations (Figure S1H). An additional, 1991 genes were only detected in WT cells (Figure S1G), including genes expressed in more than 1000 WT cells and absent from all 810 MZ*nps* cells (Figure S1K).To understand how the transcriptomes between MZ*nps* and WT donor cells compared, we reduced the dimensionality of the total dataset (selecting the top 100 principle components of expression variation for 6,632 highly variable genes) and then we embedded the dataset in two dimensions, preserving local and global data geometry (Moon *et al*., 2019). We observed that WT host cells aligned well with MZ*nps* and WT donor cells (Figure 2F), indicating that potential batch effects due to sample and library preparation were minimal. WT donor cells aligned with WT host cells (Figure 2F), indicating their transcriptome closely resembled that of host tissues. In contrast, MZ*nps* mutant cells largely segregated from both donor WT and host WT cells (Figure 2F).

We identified 825 genes significantly (FDR < 0.05, LFC > 2) upregulated in the WT cells relative to MZ*nps*, which were enriched for genes that are expressed in the first- and second-wave of ZGA, bound by NPS, and strongly downregulated in the absence of NPS at 4 hpf (Figure S1J). Conversely, the 1408 genes upregulated in MZ*nps* donor cells relative to WT cells were enriched for *miR-430* seed sequences (Figure S1J). There was also a significant overlap (151 genes, p= 4×10^-26^) between genes upregulated in MZ*nps* cells and those upregulated in MZ*nps* mutant embryos at 6 hpf, identified using publicly available bulk transcriptome data (Figure S1I) (Riesle *et al*., 2023). Importantly, 1257 genes upregulated in MZ*nps* cells at 12 hpf were not significantly upregulated in MZ*nps* embryos at 6 hpf (Figure S1I). This suggests that cells lacking NPS function during developmental reprogramming may activate distinct gene expression programs in a heterologous WT environment.

### MZ*nps* cells acquire diverse transcriptional states

The analysis of gene expression across MZ*nps* allows us to interrogate what developmental channels are established in the absence of cellular reprogramming mediated by the pioneer factors NPS after fertilization. We reasoned that MZ*nps* cells might follow one of two possible paths: i) they may collapse into a single developmental state or ii) they may acquire diverse developmental states. To investigate this, we constructed a graph of gene expression similarity between MZ*nps* cells, which was used to cluster cells into sub-populations with distinct expression characteristics (Figure 3A and 3B). We identified five sub-populations that were represented in both mutant donor replicates (KO 1-5) (Figure 3B). Differentially expressed genes enriched within each sub-population included developmental TFs typically expressed in distinct tissues during development (Figure 3C–D). KO1, the most abundant sub-population accounting for more than 60% of MZ*nps* cells (n=550), was associated with the enriched expression of *meis1a* and *anxa11a* – typically expressed in the somites and the notochord, respectively. KO2 (n=132) was enriched for expression of *ved*, a gene that acts as a ventralizing factor and transcriptional repressor to inhibit expression of organizer genes (Shimizu *et al*., 2002). KO3 (n=93) was characterized by the upregulation of *sox2* and *sox19a*, SoxB1 TFs involved in nervous system development (Okuda *et al*., 2010), as well as *otx1*, *otx2a,* and *otx2b* which are involved in brain and eye development (Martinez-Morales *et al*., 2001; Lane and Lister, 2012). GO term enrichment analyses for biological processes coincided with “generation of neurons (GO:0048699)” and “neuron differentiation (GO:0030182). KO4 similarly had GO term enrichment for neurogenesis-related terms, as it highly expressed *onecut1* and *onecutl,* which regulate differentiation and distribution of specific neural cell types (n=24) (van der Raadt *et al*., 2019). Finally, KO5 (n=11) was enriched for *nanos3, ddx4,* and *h1m*, all genes with critical roles in primordial germ cell (PGC) development. Together, these data suggest that MZ*nps* cells in a WT environment are not a homogenous population; instead, they represent molecularly distinct developmental states.

**Figure 3.**
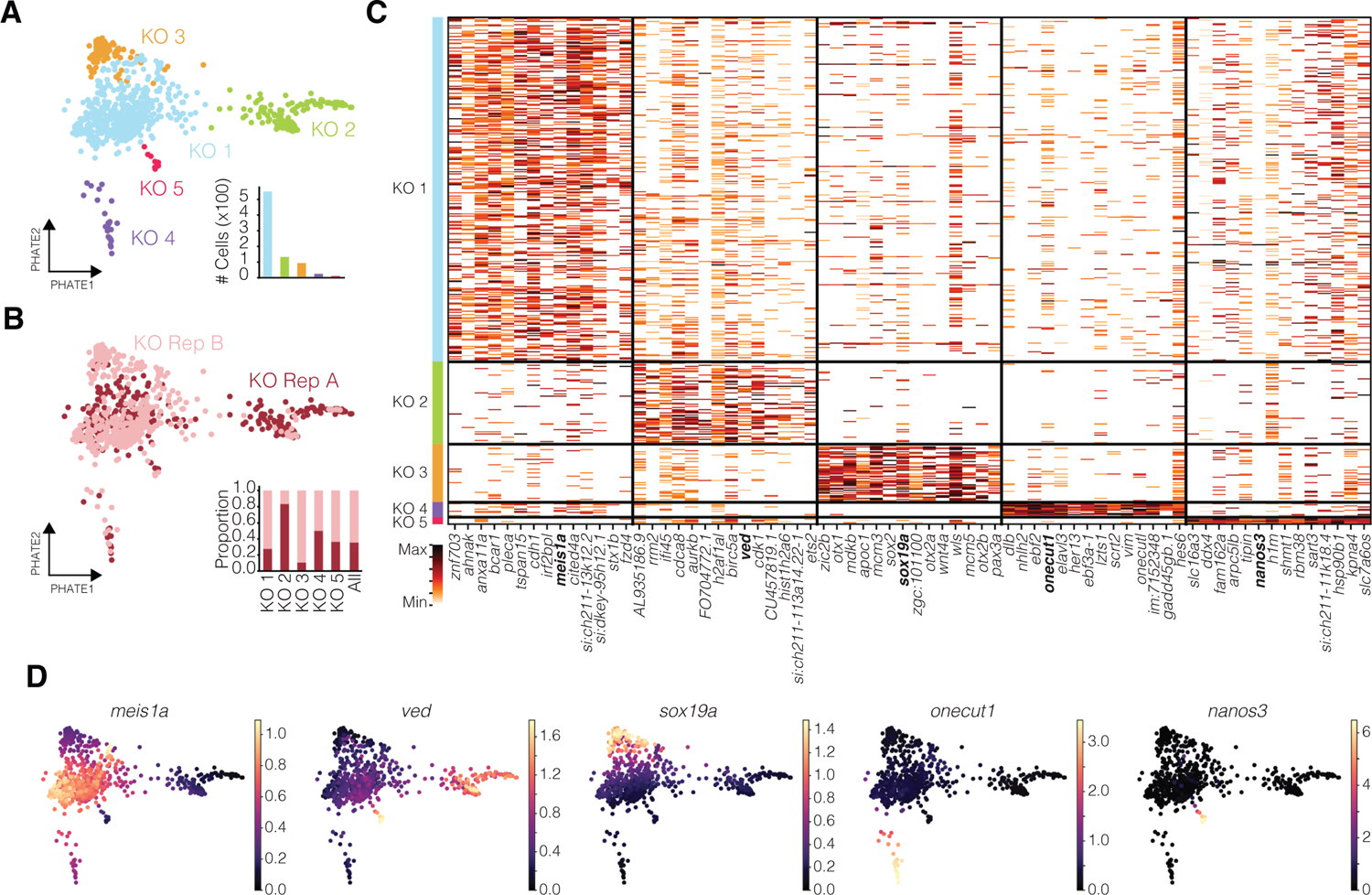
*MZnps* cells acquire distinct cell states *in vivo* **A.** 2D PHATE embedding of MZnps donor cells (810 total) colored by 5 sub-populations (KO 1-5). Bar plot shows the number of cells (×100) per sub-populations. **B.** 2D PHATE embedding colored by replicate and bar plot showing the representation of cells from each KO sub-population in two independent replicates. KO Rep A (Replicate A), maroon. KO Rep B (Replicate B), pink. **C.** Heatmap showing the normalized expression in single cells of the for top 14 marker genes in each MZnps sub-population. Rows correspond to cells, columns correspond to genes, expression is indicated by color (min-max normalized across cells). color-coded by cluster. **D.** 2D PHATE embedding colored by smoothed expression of select transcription factors identified among markers of MZ*nps* sub-populations (KO1: *meis1a,* KO2: *ved,* KO3: *sox19a,* KO4: *onecut1,* KO5: *nanos3)*.

### MZ*nps* cells resemble, but are distinct from, WT cells

We next asked whether MZ*nps* cells are canalized towards WT cellular states, or do they follow new paths representing unique developmental programs or cellular states? Because the gene expression program of a given cell depends on developmental time, we first explored whether *MZnps* cells could represent different temporal states compared to wild type cells. To estimate the developmental age of WT and MZ*nps* cells (Figure S2A; Methods), we first used publicly available reference scRNA-seq to annotate WT cell types (from Farrell *et al*., 2018; Lange *et al*., 2023; Sur *et al*., 2023), and then we compared our samples to a time series spanning 3.3-12 hpf in zebrafish (Farrell *et al*., 2018) (Figures 4A and S2A). Consistent with their time point of collection, the average predicted age of WT host and donor cells was 10.59 hpf (±0.008 CI), with some significant deviations from this mean in tissues that are specified earlier in development including PGCs, the tailbud and cells from the enveloping layer (Figure S2A).

**Figure 4.**
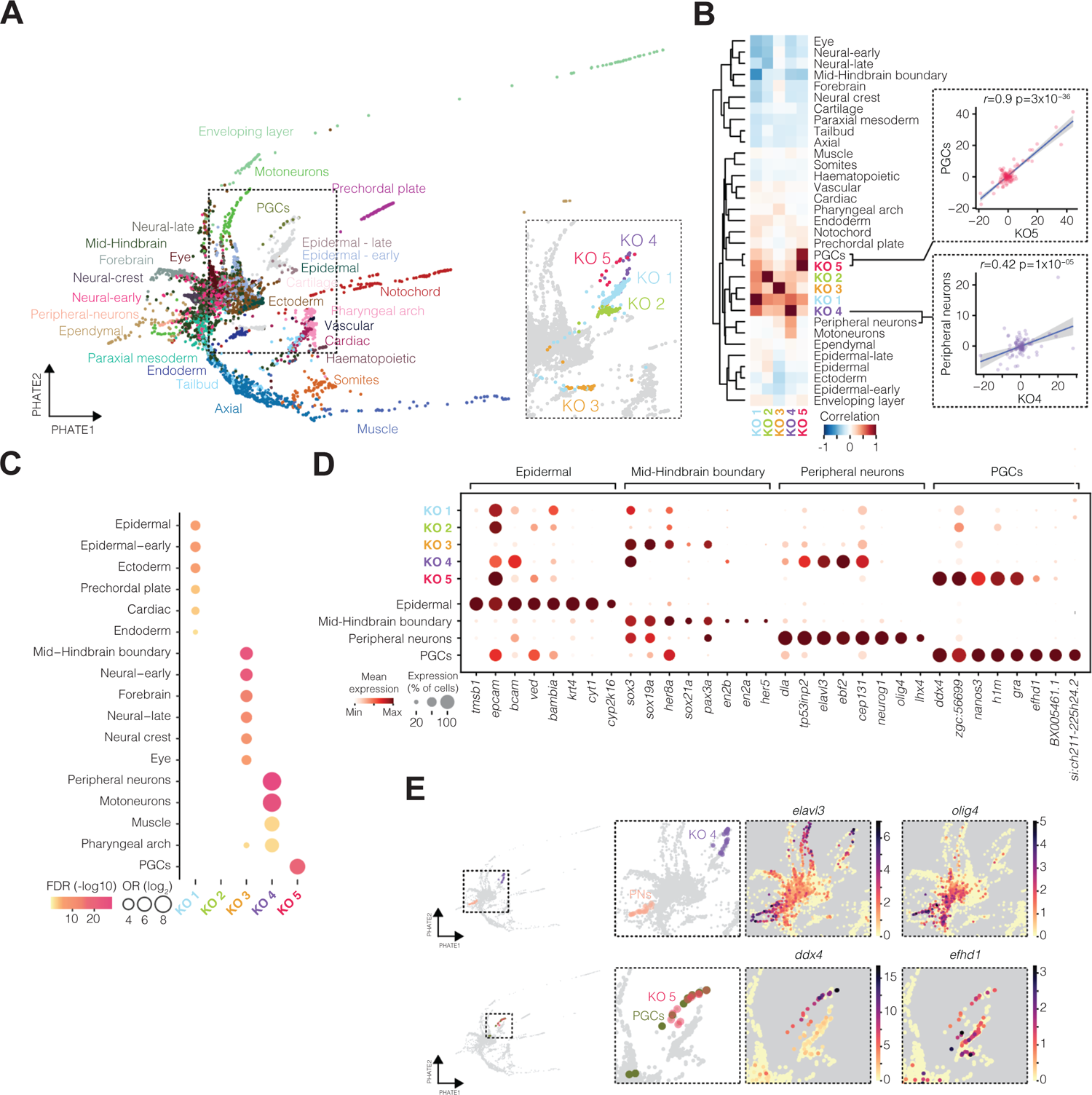
*MZnps* cell states mostly are distinct from, but correlated with, wild type cell states **A.** 2D PHATE embedding of total dataset, colored by WT cell tissue annotations (MZ*nps* cells colored grey). Inset, magnified view of MZ*nps* cells colored by KO sub-populations (WT cells colored grey). Custom annotations were generated using single-cell RNA-seq datasets from Farrell et al., 2018. **B.** Pearson correlation and hierarchical clustering (complete) between the transcriptome (top 100 PCs) of *MZnps* sub-populations (KO 1-5) and WT tissues. Top right, correlation between wild type Primordial Germ Cells (PGCs) and KO 5 (r=0.9, p=3×10^-36^). Bottom right, correlation between peripheral neurons and KO4 (r=0.42, p=1×10^-05^). Axes represent scaled expression in each tissue. **C.** Dot plot of WT tissue marker enrichment among marker genes of MZnps sub-populations. Dots indicate significant enrichment with dot size representing the odd-ratio (log_2_) and color the false discovery rate (FDR, -log_10_). **D.** Dot plot showing the mean expression (color) and detection (size, % of group) for top marker genes associated with the epidermal, midbrain-hindbrain boundary, peripheral neurons, and PGC cells, in cells of MZnps sub-populations (KO1-5) and the corresponding WT tissues. Rows correspond to cell types, columns correspond to genes, color is scaled column wise (Min-Max). **E.** 2-D PHATE embedding colored by MZnps sub-populations and correlated WT tissues and the smoothed expression of select markers of WT tissues. Top, labels for KO4 or peripheral neuron (PNs) cells, with expression of *elavl3* or *olig4.* Bottom, labels for KO5 or PGCs, with expression of *ddx4* or *efhd1*.

Similarly, the mean estimated age of MZ*nps* sub-populations KO1-4 was around 10.28 hpf (±0.037 CI) and was not different from the combined mean estimated for WT cells (Figure S2A). The estimated age of KO5 was significantly lower than the other MZnps sub-populations (∼7 hpf) yet similar to PGCs, consistent with the upregulation of PGC marker genes and with PGCs being directed down a distinct developmental canal early in development (Extavour and Akam, 2003; Farrell *et al*., 2018; Hansen and Pelegri, 2021). Together, these results suggest that KO cells represent a similar developmental stage to wild type cells and are not stalled in a preceding developmental state.

Do KO sub-populations resemble any WT cell states? First, to confirm that MZ*nps* cells represented real cell states, we compared the number of genes expressed in the MZ*nps* sub-populations to cells from WT tissues and found that they were generally comparable with WT tissues, with KO2 expressing the fewest genes (Figure S3A). Next, we assessed the similarity between WT cell-types and MZ*nps* KO sub-populations. To this end, we quantified the gene expression variance between each WT and MZ*nps* cell states. We observed a strong correlation between knockout sub-populations KO5 and KO4 with WT tissues (Figures 4B and S3A and B). KO5 was strongly correlated (rho=0.89, p=3×10^-36^) with WT primordial germ cells (PGCs), indeed stronger than its resemblance to KO cells from the other sub-populations. PGC marker genes were highly over-represented among genes upregulated in KO5 which included most of the known PGC markers such as *nanos3, ca15b, dnd1,* and *ddx4 (vasa)* (Figure 4C-4E).

Importantly, KO5 cells are the only MZ*nps* cells that are clustered together with WT cells, co-embedding with PGCs (Figure 4E). Second, the KO4 sub-population was correlated with WT peripheral-neurons (rho 0.42, p=1×10^-5^) and motoneurons (rho 0.35, p=4×10^-4^), albeit to a lesser extent than KO5 with PGCs (Figures 4B and C, and S3B). While also significantly enriched for marker genes of peripheral neurons/motoneurons, several key genes associated with neuron formation, including *olig4* and *neurog1*, were not expressed in KO4 (Figure 4D). Concordantly, KO4 cells and peripheral neurons localized to distinct areas of the embedding (Figure 4E), suggesting a moderate resemblance between these cell types. While KO1 and KO3 transcriptomes were not significantly correlated with wild type tissues overall, they did show some significant overlap in marker gene expression with non-neural ectoderm and neural tissues, respectively. KO3 was significantly enriched for genes found at the mid-hindbrain boundary (OR 19.41, FDR=6.27×10^-25^) involved in neural development, including *sox3* and *her8a* (Figure 4D). KO1 was enriched for markers of epidermal cells (OR 8.34, FDR=9.96×10^-8^) including *krt8* and *epcam*, and it also showed elevated expression of apoptosis-associated genes (95% above mean WT expression levels), suggesting that these cells may be engaged in programmed cell death (Figure S2B). Given that KO1 represents ∼60% of the cells, these results are consistent with the increase in cell death and cleaved Caspase3 expression in a subset of cells (Figure S2D and E). Thus, KO1 may represent a reprogramming impasse towards further development (developmental dead-end). In contrast, different WT cell types expressed varying levels of apoptosis gene programs, which resembled levels observed in the other KO clusters (Figure S2C). Lastly, KO2 appeared as a distinct cell state that showed no significant resemblance to WT cell-types, in terms of either transcriptome correlation or marker expression (Figure 4B-D).

Together, these results suggest that MZ*nps* cells can specify cells highly resembling germ cells, while other cellular states are distinct from those that developed in wild type embryos.

### Modes of regulation in embryonic cells in the absence of reprogramming by NPS

Cell differentiation is driven by lineage specific transcription factors that regulate gene expression modules that are refined post-transcriptionally by microRNAs and RNA binding proteins. In the Waddington framework, the upregulation of such modules direct the canalization path of cells into differentiated states. We investigated the regulatory networks shaping gene expression and cellular diversification in the absence of the reprogramming factors NPS. We identified 42 gene expression modules (EM) across WT and MZ*nps* KO sub-populations, 14 of which were enriched MZ*nps* cell subpopulations (Figure 5A; Methods). We reasoned that the generation of new cell states in the absence of the pioneer factors NPS could be derived from a defective nuclear or cytoplasmic reprogramming after fertilization, alternatively, mutant cells might be able to activate the gene expression modules present in wild type cells through compensatory or mechanisms independent of NPS. To test these hypotheses, we compared the regulation of different gene expression modules in WT and MZ*nps* embryos between 2.5 – 6 hpf, while analyzing i) the NPS binding profile, ii) the presence of *miR-430* target sites, and iii) the chromatin accessibility for each EM.

**Figure 5.**
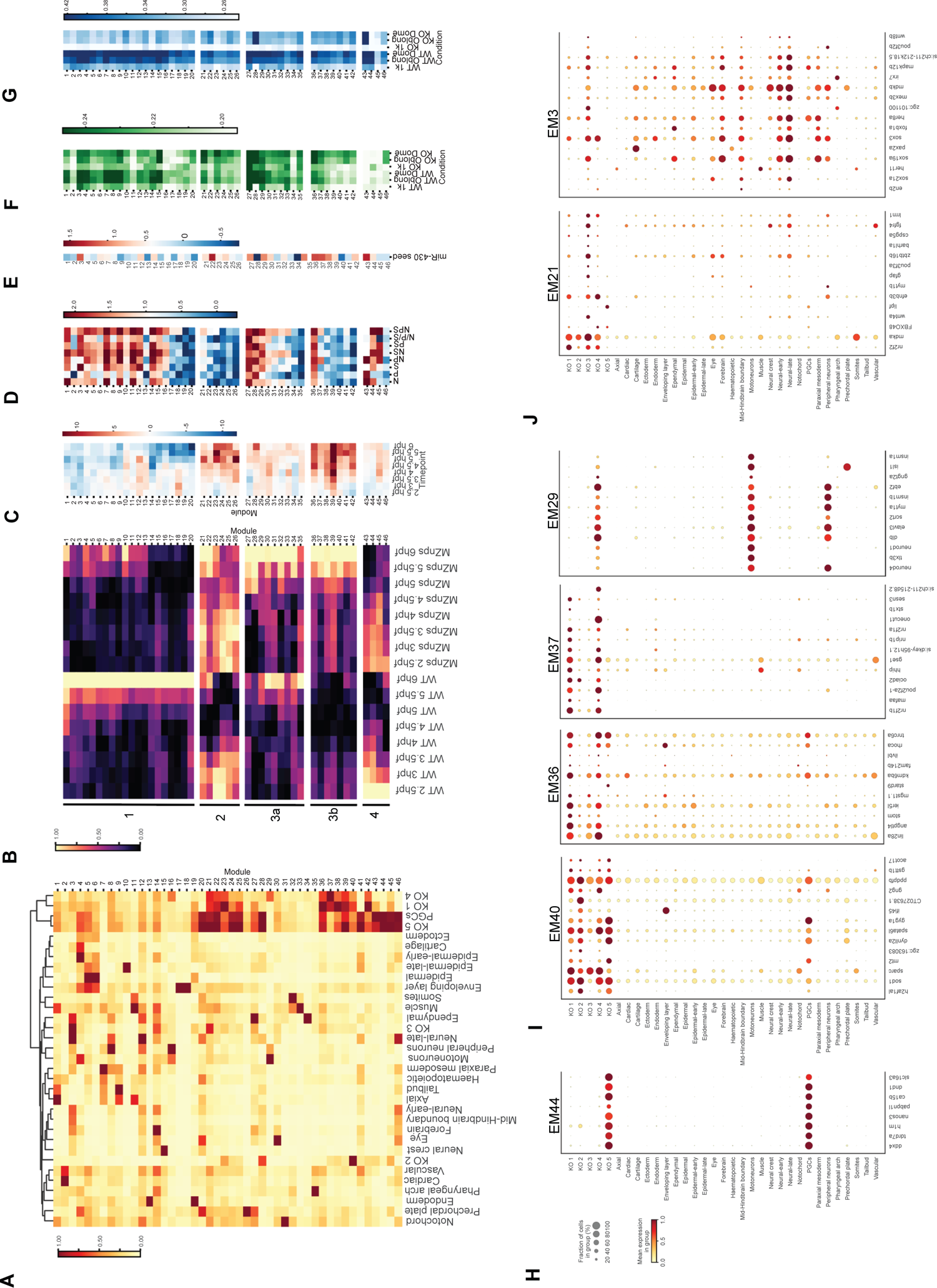
Co-expression of novel transcription factor combinations in *MZnps* cells represent novel *in vivo* cell states **A.** Heatmap of EM expression and hierarchical clustering (Ward) of WT tissues and MZnps sub-populations. Rows correspond to EMs while columns correspond to cell types. Heatmap in scaled by row (min-max). **B.** Heatmap of EM scores in WT and MZnps embryos (bulk transcriptomics) at 8 timepoints between 2.5 hours post-fertilization (hpf) and 6 hpf. Rows correspond to EMs (as identified in A), columns correspond to samples. Heatmap is scaled by row (min-max). Data from Riesle et al., 2023. **C.** T-statistics of the difference in expression of each EM in MZnps embryos relative to WT at each timepoint in B. Red represents EMs with increased expression in MZnps embryos. Data from Riesle et al., 2023. **D.** Heatmap of enrichment for transcription factor (TFs) peaks at enhancers (accessible regions within +/-5kb of transcription starts sites) among genes in each EM identified in A. Rows correspond to EMs, columns correspond to individual TFs and combinations as indicated (N: Nanog; P: Pou5f3; S: Sox19b). Color corresponds to the log odds ratio of enrichment. Data from (Xu *et al*., 2012) and (Miao et al., 2022). **E.** Heatmap of enrichment for *miR-430* seed sequences (GCACTT) within the 3’UTRs of genes in each EM identified in A. Rows correspond to EMs and color correspond to the log odds ratio of *miR-430* seed sequence enrichment. **F.** Heatmap of chromatin accessibility (mean normalized ATAC read counts) at promoters (transcription start site, TSS, +/- 500bp) of genes in each EM identified in A. Green, more open. Ordered by WT (3, 3.7, and 4.3 hpf) and *MZnps* (3, 3.7, 4.3 hpf). Data from Riesle et al., 2023. **G.** Heatmap of chromatin accessibility (mean normalized ATAC read counts) at enhancers (+/- 200bp around summit of ATAC accessible peaks within 5Kb of a TSS) of genes in each EM identified in A. Green, more open. Ordered by WT (3, 3.7, and 4.3 hpf) and *MZnps* (3, 3.7, 4.3 hpf). Data from Riesle et al., 2023. **H.** Dot plot showing the mean expression (color) and detection (size, % of group) for the top 8 genes (ranked by importance) in EM44 across WT tissues and MZ*nps* sub-populations. Rows correspond to cell types, columns correspond to genes, color is scaled column wise (Min-Max). **I.** Dot plots showing the mean expression (color) and detection (size, % of group) for the top 8 genes (ranked by importance) in EMs 40, 36, 37, and 29 across WT tissues and MZ*nps* sub-populations. Rows correspond to cell types, columns correspond to genes, color is scaled column wise (Min-Max). **J.** Dot plots showing the mean expression (color) and detection (size, % of group) for the top 8 genes (ranked by importance) in EMs 21 and 3 across WT tissues and MZ*nps* sub-populations. Rows correspond to cell types, columns correspond to genes, color is scaled column wise (Min-Max).

Interestingly, we identified four main patterns of gene regulation in these cells (Figure 5B and S4). The first pattern corresponds to a failure in nuclear reprogramming characterized by the enrichment of NPS binding, reduced chromatin accessibility and the strong downregulation of these modules (EMs 1, 4, 5, 6, 7, 10, 11, and 12) in the absence of NPS (Figure 5B-G). The second pattern corresponds to a failure in cytoplasmic reprogramming due to the loss of *miR-430-*dependent mRNA clearance (EMs 21-26). This failure is due to a decrease in miR-430 expression, which itself requires NPS function. In the absence of NPS, these genes are characterized by strong maternal perdurance and increased zygotic expression (Figure 5B-G).

Additionally, their regulatory sequences show a depletion of NPS binding and are enriched for miR-430 target sites (Figure 5E). The third pattern corresponds to genes that undergo strong zygotic upregulation in *MZnps* mutants. These genes are newly activated by factors that either require NPS for repression as part of nuclear reprogramming (3a) (EMs 27-35), or that compete with NPS for the transcriptional machinery as part of a compensatory mechanism (3b) (EMs 36-42) (Figure 5B-G). They are characterized generally by an enrichment of NPS binding (3a) or a depletion of NPS binding (3b), respectively (Figure 5D). The last pattern corresponds to those specifically defining PGCs and KO5 (EMs 43-46), which include genes such as *ddx4, dnd1,* and *nanos3* and their expression was generally unaffected by the loss of NPS (Figure 5B-D).

Altogether, these results suggest that the different gene expression programs channeled in the absence of NPS pioneering factors, derive from failures in nuclear and cytoplasmic reprogramming as well as compensatory gene regulation.

### Developmental states defined in the absence of NPS-mediated reprogramming

Finally, we investigated what cell states (defined by specific EM combinations) were acquired in MZ*nps* cells. Most WT tissues were defined by the specific enrichment of a small number of EMs (for example, EM15 for neural crest, EM17 for enveloping layer, and EM11 for mesodermally-derived lineages) (Figure 5A). These WT-enriched modules showed reduced expression in MZ*nps* sub-populations, suggesting NPS-mediated reprogramming precludes the canalization towards these WT cell states.

Next, we explored whether MZ*nps* cells canalize towards known WT states. Consistent with previous analyses (Figure 4B), canalization to PGCs was largely unaffected in MZ*nps* cells, as there was a strong resemblance uniquely between KO5 and WT PGCs (Figure 5H). These data suggest that many of the gene expression programs that define PGCs are independent of NPS-reprogramming. Interestingly, sub-populations KO1 and KO4 were also enriched for multiple EM shared with KO5 and PGCs, suggesting a mixed identity of those cells.

We next considered two cell groups that may represent related subpopulations along a novel canalization attempt (K1 and KO4). KO1 was enriched for multiple modules that are strongly activated in the absence of reprogramming by NPS (Figure 5A and B) and are associated with cell-stress (including heat-shock factors in EM3) (Figure 5I). This suggests these programs related to cell stress may be independent of NPS. These canalization attempts can result in a mixed identity, such as the one observed in the KO4, which has EMs shared with KO1-5 (EM40), KO1 and 3 (EM36), and just KO1 (EM37) (Figure 5I). Interestingly, the top gene expressed in EM36 was *lin28a,* an RNA-binding protein shown to inhibit differentiation states as it helps during *in vitro* reprogramming to IPSCs (Figure 5I) (Melton et al., 2010; Wang et al., 2019; Zhang et al., 2016). Thus, EM36 may represent a gene expression module that compensates for the absence of NPS in somatic cells. This mixed identity is also supported by the co-expression of *neurod1/4*, *tlx3b*, *elavl3* (EM29), which is only shared between KO4, motoneurons and peripheral neurons (Figure 5I). Together, KO1 may represent a cell state in an inappropriate environment, while KO4 may represent a receptive cell state in a permissive environment canalizing towards a mechanosensory neuron-like state.

Another group of cells presented a mixed state (KO3) manifested by the combined expression of only two EMs: one enriched in KO cells (EM21) and another enriched in WT neural tissues such as late neural progenitors and ependymal cells (including *dmbx2*, *her8a*, and Sox TFs *sox21a, sox3*, and *sox19a)* (EM3) (Figure 5J). Thus, we propose that KO3 represents a novel cellular state not present in early embryogenesis characterized by the integrated expression programs from WT neural lineages with novel modules induced in the absence of NPS. Together, these data support that MZ*nps* cells, which lack the first nuclear and cytoplasmic reprogramming events, can create new developmental “canals” *in vivo*.

## Discussion

The maternal-to-zygotic transition (MZT) provides a developmental framework to understand the implications of the nuclear and cytoplasmic reprogramming from the differentiated state of the sperm and the oocyte into a transient totipotent state (Yartseva and Giraldez, 2015). Here, we demonstrate that cells with mutations in the pioneer factors NPS are defective in both cytoplasmic and nuclear reprogramming and fail to acquire differentiated wild-type states. We observed that some MZ*nps* cells likely reach a developmental dead-end, while others achieve distinct mixed fates with multiple expression modules present, indicating that cells lacking reprogramming factors can adopt new cell states *in vivo*. Interestingly, we found that multiple MZ*nps* cells closely resemble primordial germ cells, suggesting that their development does not require major reprogramming by NPS. These findings provide novel insights to understand how developmental reprogramming by pioneer factors establishes the Waddington epigenetic landscape to canalize cell differentiation during development.

In the developmental reprogramming hypothesis, both nuclear and cytoplasm reprogramming are required for this successful maternal-to-zygotic transition to enable correct cell differentiation (Giraldez, 2010; Yartseva and Giraldez, 2015). Indeed, the process of developmental reprogramming was first shown through nuclear transfer experiments, which elegantly demonstrated that the oocyte’s cytoplasm has the information required to reprogram a differentiated cell’s nucleus into a transient totipotent state that can then achieve multiple cell fates *in vivo* (Gurdon, Elsdale and Fischberg, 1958). In zebrafish, the oocyte provides maternal pioneer factors – Nanog, Pou5f3/OCT4, and Sox19b/SOX2 (NPS) – that are required for the initial nuclear reprogramming and subsequent cytoplasmic reprogramming through the NPS-dependent activation of *miR-430* (Lee *et al*., 2013; Miao *et al*., 2022). The effective reprogramming during the MZT is proposed to influence the subsequent landscape of cell differentiation, however testing this model *in vivo* has remained challenging given the gastrulation defects in MZ*nps* mutant embryos. To overcome this challenge, we used cell transplantation to allow the developmental progression of MZ*nps* mutant cells. This approach allowed us to interrogate whether cells lacking nuclear and cytoplasmic reprogramming adhere to a singular, homogenous state, or if they are directed towards wild type or novel developmental states. We find that mutant cells for NPS can indeed adopt novel cell states and gene expression modules distinct from WT. The MZ*nps* sub-populations of KO1-4 cells exhibited unique transcriptional profiles resembling cell states not observed in developing wild type cells, with sub-populations KO4 and, to a lesser extent, KO3 showing similarities to peripheral and central neural fates, aligning with a neural default model (Muñoz-Sanjuán and Brivanlou, 2002). Other cell clusters displayed novel somatic signatures likely established early in development. The exception was the KO5 MZ*nps* cells, which highly resembled WT primordial germ cells (PGCs). These results suggest that Nanog, Pou5f3, and Sox19b may be dispensable for PGC gene regulatory programs and lineage specification. We propose that MZ*nps* mutant cells can forge new ‘canals’ along their developmental trajectories through *de novo* combinations of gene expression modules not observed in wild type cell states.

Our results support that the initial reprogramming steps lay the groundwork for later gene expression programs. Nanog, Pou5f3, and Sox19b are pioneer factors that directly regulate chromatin accessibility for successful zygotic genome activation and subsequent clearance of the maternal program by activating *miR-430* (Miao et al., 2022). Yet, the nuclear reprogramming by NPS is not only required for genome activation, since other genes still get activated in the absence of NPS (Onichtchouk *et al*., 2010; Miao *et al*., 2022). We find that some EMs (EM3) can bypass NPS function later in development, despite requiring NPS-dependent reprogramming and direct NPS binding during ZGA. Interestingly, other EMs were upregulated in the absence of NPS (for example, EM grouping 3b). These results suggest that NPS plays not only a direct role in ZGA, but also an indirect role likely through the competition with other transcription factors for the transcriptional machinery early in development. Future studies will be needed to delineate this dual role of NPS, which underscores its importance not just in activating genome-wide transcription but also through a likely competitive and cooperative mechanism with other transcription factors early in development.

Using the Waddington landscape as a framework, our results contrast to previously published findings, which suggest that mutant or perturbed cells typically follow established wild type developmental paths and do not forge new ones. For example, cells lacking the coreceptor for the mesendoderm inducer Nodal show reduced expression of relevant gene modules but do not generate novel mutant-specific modules; instead, these mutant cells are canalized into a subset of wild type fates (Farrell *et al*., 2018).These results are consistent with the elimination of one of the canals in the developmental landscape (mesendoderm), and thus cells take the other canals available for development. Our data suggest that even in the absence of successful reprogramming, cells can still navigate developmental decisions where distinct combinations of genes and transcription factors can canalize cells toward new developmental states. We propose that the failure to reprogram may result in either “dead-end” states, similar to those seen in failed nuclear transfer or induced reprogramming attempts (Gurdon, Elsdale and Fischberg, 1958; Biddy *et al*., 2018), or alternatively novel “mixed” states. Conceptually, these new states may represent the creation of new canals along with the simultaneous failure to irreversibly close off other canals (Ferrell, 2012).

Altogether, this study shows how NPS-dependent nuclear and cellular reprogramming sculpt the developmental landscape *in vivo*, essential for guiding the precise developmental decisions that underpin differentiation. This study not only refines our understanding of the role of pioneering factors in developmental reprogramming, but also illustrates how variations in early reprogramming can reshape the Waddington landscape, leading to unexpected and diverse cell fates.

---

**Figure S1.**
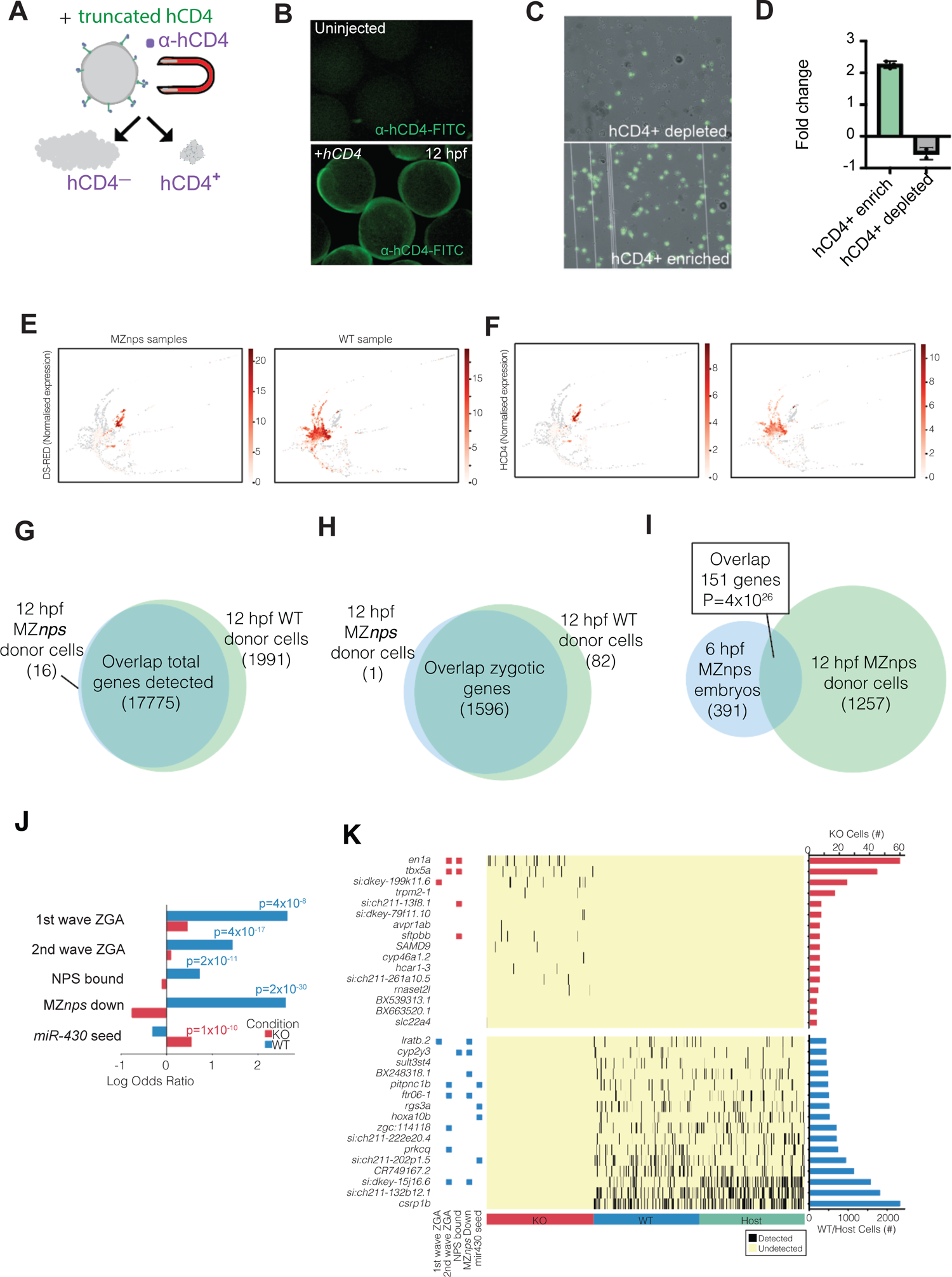
Isolation and identification of labeled *MZnps* donor cells. Related to Figures 1 and 2. **A.** Schematic of magnetic enrichment for hCD4+ cells (cyan) using anti-hCD4 MicroBeads following dissociation of embryos at 12 hpf. Left, hCD4-depleted population. Right, hCD4+ enriched population. **B.** Wild type zebrafish embryos uninjected (top) or injected with hCD4-*prrg2-3’UTR* mRNA (bottom) at the one-cell stage, stained with α-hCD4-FITC (green) at 12 hpf. **C.** Representative image of magnetic enrichment for hCD4+ cells (cyan) using anti-hCD4 MicroBeads following dissociation of embryos at 12 hpf. α-hCD4-FITC staining (green) in an enriched sample (right) and depleted sample (left). Right, CD4+ enriched population. **D.** Fold change in the proportion of labelled cells of hCD4+ cells relative to expected cell proportions. *Tg(H2B-GFP)* embryos were injected with hCD4-*prrg2-3’UTR* mRNA at the one-cell stage. 8 injected *Tg(H2B-GFP)* embryos were mixed with 25 wild type embryos, dissociated, and enriched using CD4 MicroBeads at 11 hpf. **E.** 2-D PHATE embedding of total datasets colored by normalized *DsRed* mRNA expression in cells from KO donor (left, KO and Host in grey) and KO (right, WT and Host in grey). **F.** 2-D PHATE embedding of total datasets colored by normalized *hCD4* mRNA expression in cells from KO donor (left, KO and Host in grey) and KO (right, WT and Host in grey). **G.** Venn diagram showing number and overlap of genes detected in MZ*nps* or WT cells. **H.** Venn diagram showing number and overlap of strictly zygotically expressed genes detected in MZ*nps* or WT cells. **I.** Venn diagram showing number and overlap of genes upregulated in MZ*nps* cells relative to WT cells at 12hpf and those upregulated in MZ*nps* embryos relative to WT at 6hpf. **J.** Enrichment of genes sets associated with the first wave ZGA, second wave ZGA, NPS bound, downregulated and mir430-seed containing 3’UTRs among genes differentially expressed (FDR < 0.05, LFC < −2: blue, LFC > 2: red). **K.** Heatmap showing the detection of genes in MZ*nps* (KO, red), WT (Blue) and Host (Green) cells. Black like indicate detection is a cell (UMI >1). Shown are the differentially detected genes between MZ*nps* and WT cells (top 16 most detected). Tile plots indicate their respective annotation among the genesets listed in J. Bar plots show the total detection in KO (red bars) or WT (host and donor) cells.

**Figure S2.**
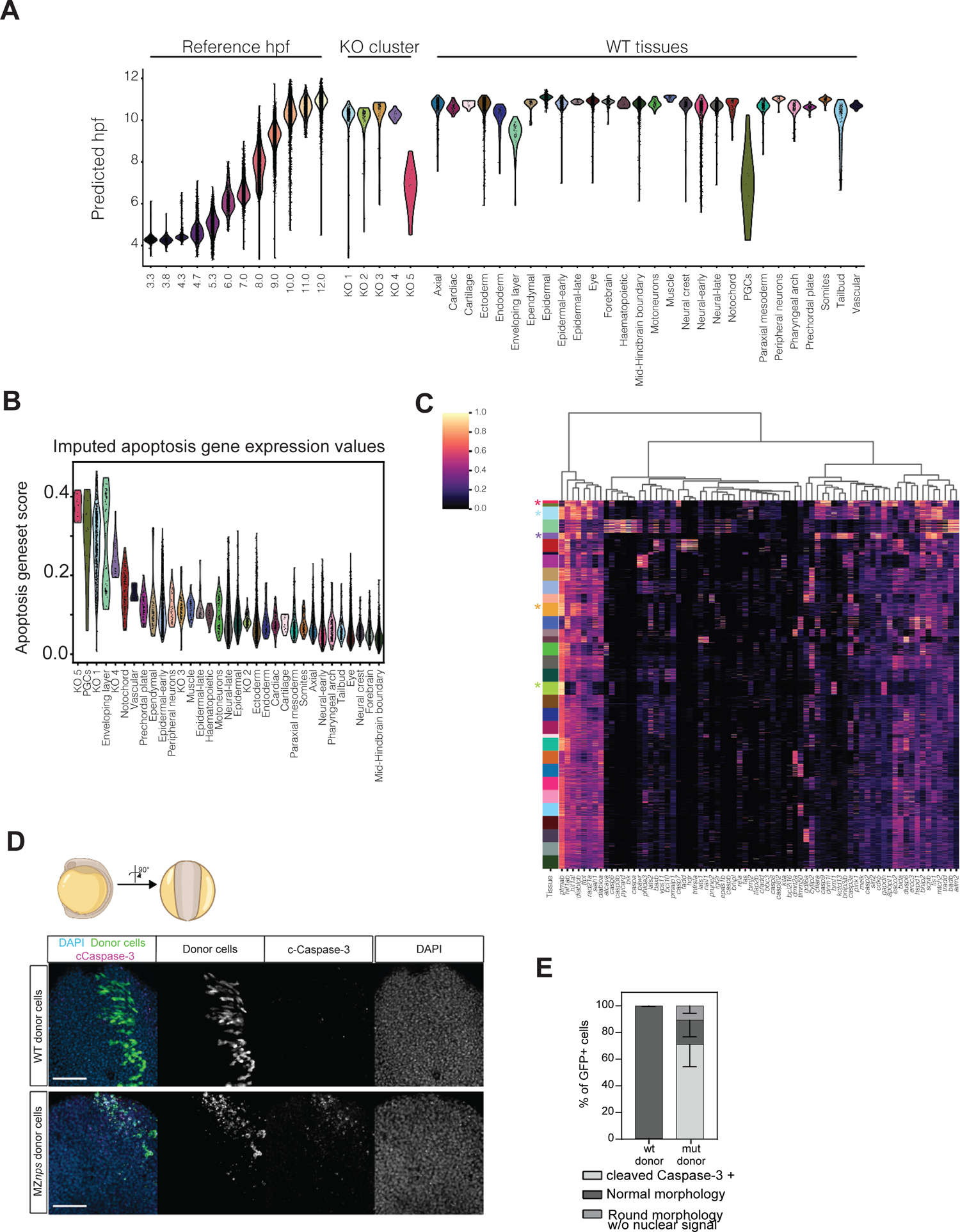
Comparisons between WT and MZ*nps* cell groups. Related to Figure 4. A. Violin plots of predicted time (in hours post-fertilization, hpf) of reference single-cell RNA-sequencing timecourse (from Farrell et al., 2018), each KO cluster, and each WT cell cluster. B. Violin plots showing the apoptosis geneses score in cells from MZnps sub-populations (KO1-5), and WT tissues. Dots represent individual cells. C. Heatmap of gene expression (smoothed) of a curated apoptosis-associated geneset. Row colors and order correspond to panel B. Asterix indicate KO sub-populations. D. WT host embryos imaged at 3-6ss containing GFP+ donor cells (green) from either WT (top) or MZ*nps* mutant embryos (bottom) that were transplanted at the 1-cell stage. Embryos are oriented dorsally. Cleaved Caspase-3 (magenta) and nuclei are labeled with DAPI (blue). Scale bar = 100μm. These embryos are also presented in Figure 1B. E. The mean percent of donor cells with normal morphology, cleaved Caspase-3 co-localization, or round morphology without nuclear signal (DAPI staining). Error bars represent standard deviation, n=3 embryos for WT donor cells and n=5 embryos for MZ*nps* donor cells.

**Figure S3.**
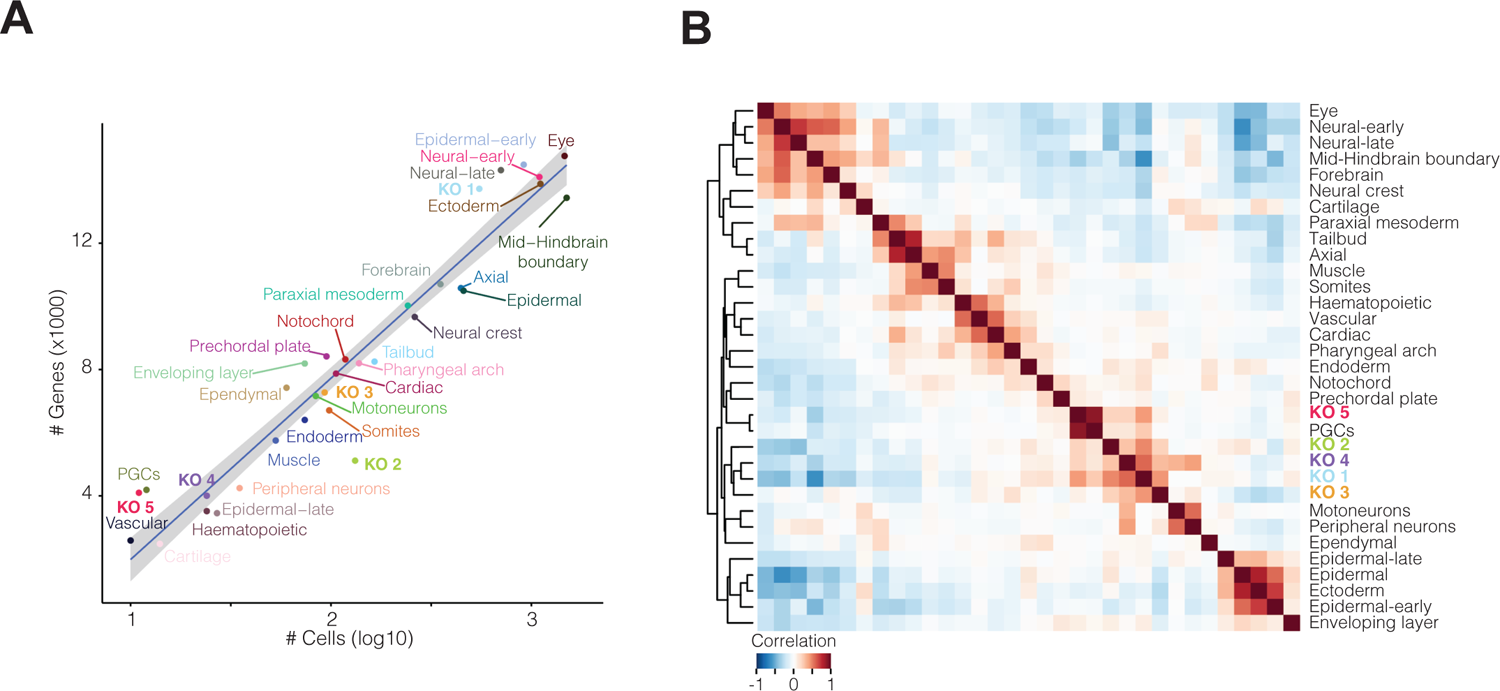
Library complexity and transcriptome similarity between MZnps and WT cells. Related to Figure 4. A. Relationship between number of cells (log_10_) and number of genes (×1000) for WT and KO cells assigned cell-type labels. Inset, number of genes (×1000) expressed in individual MZ*nps* (KO) clusters. B. Pearson correlation and hierarchical clustering (complete) between the transcriptome (top 100 PCs) of *M*Z*nps* sub-populations (KO 1-5) and WT tissues. (A subset of this figure is present Figure 4B.)

**Figure S4.**
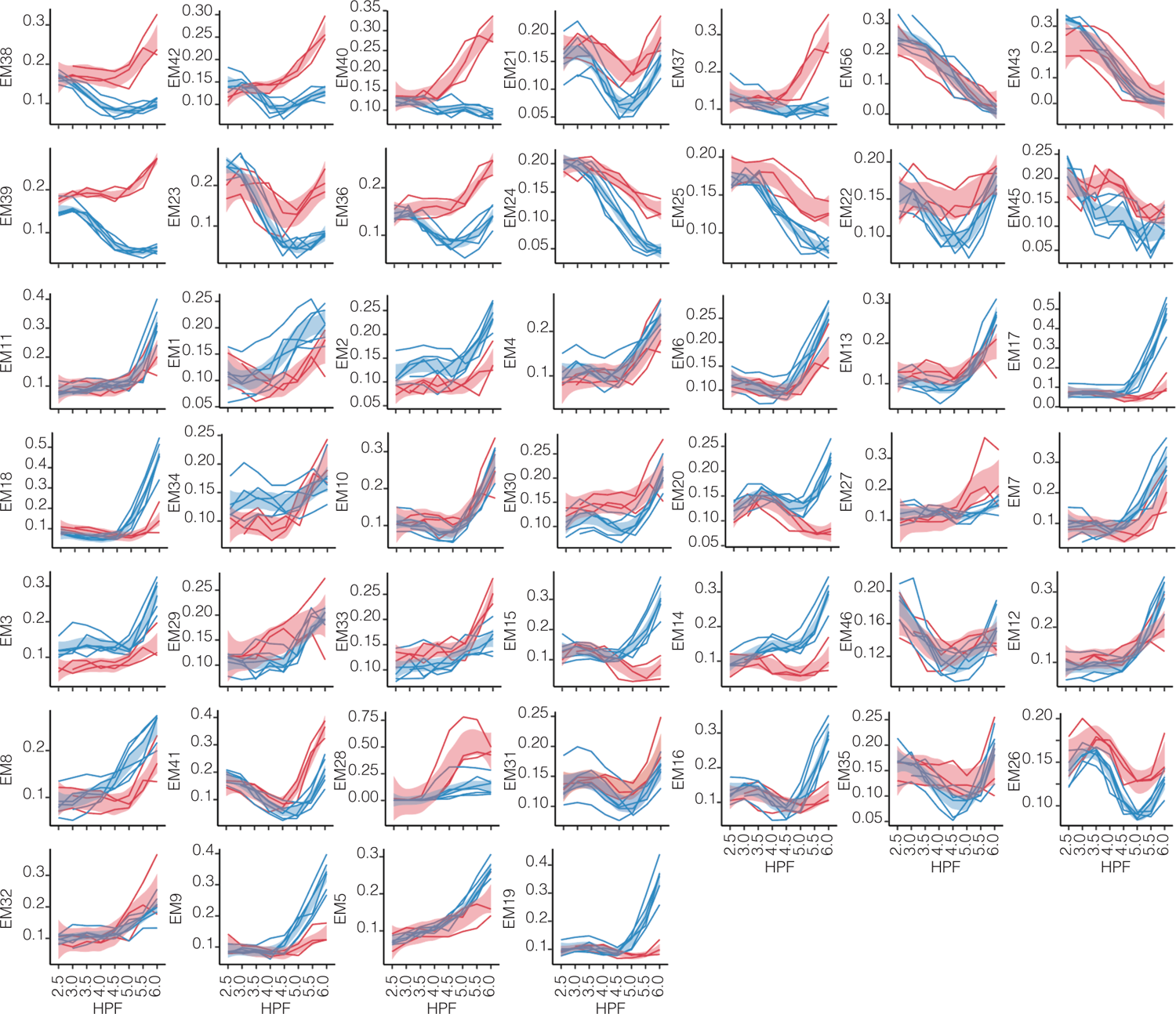
EM expression over time in KO and WT embryos. Related to Figure 5. Line plots of EM scores in WT and MZ*nps* embryos (bulk transcriptomics) at 8 timepoints between 2.5 hours post-fertilization (hpf) and 6 hpf. Each plot corresponds an EM as identified in Figure 5A, lines correspond to independent replicates colored by condition (red = MZ*nps* embryos, blue = WT embryos). Data from Riesle et al., 2023.

## Acknowledgments

We thank Guilin Wang from the Yale Center for Genome Analysis for sequencing support; Hiba Codore and Nitya Khatri for technical help; and Sarah Dube, Tim Gerson, and Damilola Olowookere for excellent animal care. This work is supported by funding from the US NIH *Eunice Kennedy Shriver* Institute for Child Health and Human Development (K99HD105001) to V.A.T; the Surdna Foundation and the Yale Genetics Venture Fund to L.M.; Human Frontiers Postdoctoral Fellowship LT0073/2022-L and EMBO Long-Term Postdoctoral Fellowship ALTF #794-2021 to C.H.; a Canadian Institutes of Health Research (CIHR) postdoctoral fellowship to C.W.B.; and R01HD100035 (NIH) to S.K. and A.J.G. and R35GM122580 (NIH) to A.J.G. The content is solely the responsibility of the authors and does not necessarily represent the official views of the National Institutes of Health or any funding sources.

## Author Contributions

L.M., V.A.T., and A.J.G. conceived this project. L.M., V.A.T., and A.J.G performed experiments. V.A.T. designed the cell isolation method and optimizations. S.E.Y. designed and performed data analyses, with L.M. and V.A.T. contributing to guiding analyses and performing *in vivo* validations. C.H. performed confocal image acquisition and analyses. C.B generated a custom annotation of cell types in previously published wild type single cell datasets (from Farrell et al., 2018). Equal contribution from D.M and M.A. for identification of an endogenous stabilizing 3’UTR. S.K and A.J.G. contributed to funding acquisition and project supervision. V.A.T. and S.E.Y. wrote the original draft, with L.M. and A.J.G. contributing to editing and revisions. All authors revised this manuscript. S.E.Y. and L.M. contributed equally to this work.

## Methods

### Zebrafish husbandry and maintenance

The husbandry of zebrafish was conducted as previously documented (Howe et al., 2016; Westerfield, 2000). The maintenance of fish lines was carried out in accordance with the AAALAC research guidelines following a protocol approved by the Yale University IACUC. The maternal-zygotic (MZ) *nanog*^-/-^;*pou5f3*^-/-^;*sox19b*^-/-^ triple homozygous mutants (MZ*nps*) were maintained as described (Miao et al., 2022). Briefly, MZ*nps* or MZ*nanog*^-/-^;*pou5f3*^+/-^;*sox19b*^-/-^ females were crossed with MZ*nps* males, and the resulting embryos were rescued with mRNA injection (25 pg *nanog* mRNA + 30 pg *pou5f3* mRNA) at the 1-cell stage. Alternatively, MZ*nanog*^+/-^;*pou5f3*^+/-^;*sox19b*^+/-^ females were crossed with MZ*nps* males, and the resulting embryos were rescued by injecting 30 pg *pou5f3* mRNA at the 1-cell stage. Progenies from MZ*nanog*^-/-^;*pou5f3*^+/-^;*sox19b*^-/-^ or MZ*nanog*^+/-^;*pou5f3*^+/-^;*sox19b*^+/-^ females were genotyped as previously described (Miao *et al*., 2022).

### Embryo injections, blastula transplants, and dissociations

Embryos were dechorionated at the 1-cell stage. 100 pg hCD4 mRNA (described below) and 50 pg DsRed capped mRNA were injected into donor embryos (both WT and MZ*nps* embryos) at the 1-cell stage. 75 pg EGFP-CAAX capped mRNA was injected into host wild-type embryos at the 1-cell stage. At 3-4 hpf, about 50 cells were taken from donor embryos and transplanted into host WT embryos. Transplantation was carried out as described (Kemp, Carmany-Rampey and Moens, 2009). Transplanted embryos were kept at 28°C until dissociation. At ∼3-6ss stage (corresponding to ∼11-12 hpf), embryos were rapidly deyolked manually with forceps in fresh, ice-cold Ringer’s solution (per 50 ml: 5.8ml 1M NaCl; 73μl 2M KCl; 250μl 1M HEPES in 43.5 ml RNase-free water). After deyolking, embryos were rinsed and gently mechanically dissociated using polished glass pipettes in fresh ice-cold Ringer’s solution, centrifuged at 4°C at 300 rpm for 3 minutes, supernatant removed, and the remaining cells were resuspended in fresh ice-cold PBE solution (1X RNase-free PBS, 0.5% BSA, 2mM EDTA). Cells were the incubated with a 1:100 hCD4 Microbeads solution (in PBS) (Miltenyi Biotec), incubated for 10 minutes on ice, and washed 2X with ice-cold PBE solution. Cells were resuspended in 1ml PBE and placed through a MACS magnetic column (MiniMACS Separator, Miltenyi Biotec). Cells were collected into a clean RNase-free Lo-Bind tube, washed 2X with ice-cold PBE, and immediately prepared for loading onto a 10X Genomics chip (v3). The protocol was optimized and enrichment efficiencies calculated using an anti-hCD4-FITC antibody (to detect expression in zebrafish embryos), and hCD4 mRNA-injected *Tg(his:GFP)* embryos, mixed 1:9 with WT embryos, and cells counted using a hemocytometer and Trypan Blue to assess cell viability.

From single cell RNA sequencing experiments, we noted an over-representation of cell types of ectodermal origin in WT donor cells relative to WT host cell populations, although all tissue types were represented, indicating a potential technical bias introduced at the site of transplantation (as shown in Figures 2E and 4A) (Kimmel, Warga and Schilling, 1990).

### hCD4 magnetic microbeads-mediated cell enrichment of transplanted cells

A truncated human CD4 sequence (corresponding to the sequence identified by hCD4 MicroBeads, Miltenyi Biotec) was cloned into a pCS2+ vector containing an SP6 promoter, a stabilizing 3’UTR derived from the zebrafish *prrg2* endogenous gene (Boswell *et al*., 2023), and an SV40 pA.

The truncated hCD4 sequence was

5’-ATGAACCGGGGAGTCCCTTTTAGGCACTTGCTTCTGGTGCTGCAACTGGCGCTCCTCCCAGCAGCCACTCAGGGAAAGAAAGTGGTGCTGGGCAAAAAAGGGGATACAGTGGAACTGACCTGTACAGCTTCCCAGAAGAAGAGCATACAATTCCACTGGAAAAACTCCAACCAGATAAAGATTCTGGGAAATCAGGGCTCCTTCTTAACTAAAGGTCCATCCAAGCTGAATGATCGCGCTGACTCAAGAAGAAGCCTTTGGGACCAAGGAAACTTCCCCCTGATCATCAAGAATCTTAAGATAGAAGACTCAGATACTTACATCTGTGAAGTGGAGGACCAGAAGGAGGAGGTGCAATTGCTAGTGTTCGGATTGACTGCCAACTCTGACACCCACCTGCTTCAGGGGCAGAGCCTGACCCTGACCTTGGAGAGCCCCCCTGGTAGTAGCCCCTCAGTGCAATGTAGGAGTCCAAGGGGTAAAAACATACAGGGGGGGAAGACCCTCTCCGTGTCTCAGCTGGAGCTCCAGGATAGTGGCACCTGGACATGCACTGTCTTGCAGAACCAGAAGAAGGTGGAGTTCAAAATAGACATCGTGGTGCTAGCTTTCCAGAAGGCCTCCAGCATAGTCTATAAGAAAGAGGGGGAACAGGTGGAGTTCTCCTTCCCACTCGCCTTTACAGTTGAAAAGCTGACGGGCAGTGGCGAGCTGTGGTGGCAGGCGGAGAGGGCTTCCTCCTCCAAGTCTTGGATCACCTTTGACCTGAAGAACAAGGAAGTGTCTGTAAAACGGGTTACCCAGGACCCTAAGCTCCAGATGGGCAAGAAGCTCCCGCTCCACCTCACCCTGCCCCAGGCCTTGCCTCAGTATGCTGGCTCTGGAAACCTCACCCTGGCCCTTGAAGCGAAAACAGGAAAGTTGCATCAGGAAGTGAACCTGGTGGTGATGAGAGCCACTCAGCTCCAGAAAAATTTGACCTGTGAGGTGTGGGGACCCACCTCCCCTAAGCTGATGCTGAGCTTGAAACTGGAGAACAAGGAGGCAAAGGTCTCGAAGCGGGAGAAGGCGGTGTGGGTGCTGAACCCTGAGGCGGGGATGTGGCAGTGTCTGCTGAGTGACTCGGGACAGGTCCT GCTGGAATCCAACATCAAGGTTCTGCCCACATGGTCGACCCCGGTGCAGCCAATGGCCCTGATTGTGCTGGGGGGCGTCGCCGGCCTCCTGCTTTTCATTGGGCTAGGCATCTTCTTCTGTGTCAGGTGCCGGCAC – 3’

### Embryo antibody staining and image acquisition for blastula transplants

Embryos were collected at ∼12 hpf, fixed overnight in cold 4% paraformaldehyde (PFA) in 1x PBS overnight, washed with 1X PBS-Tw (0.1% Tween-20) three times, permeabilized with 10μg/ml Proteinase K for 1 minute, washed with PBS-Tw three times, re-fixed with 4% PFA/PBS-Tw, and washed again with PBS-Tw three times. Embryos were blocked with 10% normal goat serum (Thermo Fisher Scientific, 50062Z) for one hour rotating at room temperature, stained with primary antibodies against GFP Tag Monoclonal Antibody (3E6) (1:500, mouse, Thermo Fisher Scientific, A11120) and cleaved Caspase-3 (rabbit, 1:500, CST 9661S) rotating overnight at 4C, washed with PBS-Tw three times at room temperature, then stained with AlexaFluor 488 anti-mouse, AlexaFluor 546 anti-rabbit, and DAPI for one hour rotating at room temperature. Embryos were washed with PBS-Tw three times. Embryos were mounted in 0.8% low melt agarose (GPG/LMP AmericanBio, AB00981-00050) in water against a no. 1.5 cover slip. Whole embryo images at 12 hpf were acquired using an upright Zeiss LSM 980 confocal microscope with an Airyscan 2 detector and an air EC Plan-Neofluar 10x/0.3 M27 objective. In detail, images were obtained at 16 Bit with 2x line averaging, bidirectional scanning, LSM scan speed 6 (pixel dwell time 0.51 μs), 1.1x optical zoom, GaAsP-PMT detectors, an image size of 4084×4084 pixels corresponding to 769.13 μm ×769.13 μm and three-dimensional optical stacks were acquired at 2 μm spacing. Samples were illuminated with the 405nm laser line (DAPI) at 0.2-1% laser power, 820V (gain) and 0 (offset); 488nm laser line (Alexa Fluor 488) at 4% laser power, 850V (gain) and 0 (offset); 561nm laser line (Alexa Fluor 546) at 1.1% laser power, 850V (gain) and 0 (offset). Raw images were deconvolved using the Airyscan software and images shown in the result section are maximum projected, with brightness and contrast adjusted separately between experiments.

Regions of interest were imaged using the above confocal scope with a LD LCI Plan-Apochromat 40x/1.2 Imm Corr DIC M2 objective with water immersion. Settings that differ from the above whole embryo imaging were as follows: LSM scan speed 7 (pixel dwell time 0.35 μs) and three-dimensional optical stacks were acquired at 0.3 μm spacing. Samples were illuminated with the 405nm laser line (DAPI) at 0.2% laser power, 820V (gain) and 0 (offset); 488nm laser line (Alexa Fluor 488) at 1% laser power, 850V (gain) and 0 (offset); 561nm laser line (Alexa Fluor 546) at 1.1% laser power, 850V (gain and 0 (offset). Images were acquired at 4084×4084 or 3016×2160-3016 pixels corresponding to 176.26 μm x 176.26 μm and 124.29 μm x 89.01-124.29 μm, respectively. The optical zoom (1.2 or 1.7) and therefore image sizes were varied to optimally image specific tissue regions. Raw images were deconvolved using the Airyscan software and then processed with Image-J software (Schneider, Rasband and Eliceiri, 2012). Images shown in the result section show maximum projected sub-stacks with brightness and contrast adjusted separately between images to optimize visualization of GFP+ donor cells in their tissue environment.

### Imaging quantification for single transplants

3D image analysis was performed in the IMARIS software (Bitplane, Oxford Instruments, Concord MA; Version: 10.0). An analysis region was defined for each image to include the embryo tissue and exclude the yolk area so that GFP+ donor cells outside this region were excluded from quantification. GFP+ cells were identified using the ‘spot’ function and a constant size of 8 μm spheres, to exclude GFP+ cell remnants. Cells were categorized based their DAPI and cleaved Caspase-3 fluorescence intensity and then manually checked.

### Data alignment and preprocessing

Single cell data was aligned to the zebrafish genome (grcz11) with the addition of sequences of GFP, DsRed and hCD4, and UMIs for each gene (annotation transcriptome plus GFP, DsRed and hCD4) quantified per cell using the CellRanger pipeline (v3.0.2, 10x Genomics). The counts matrix was loaded into python (v3.11.5) as an anndata object (v0.9.1, (Virshup *et al*., 2023) and preprocessed using scanpy (v1.9.3, (Wolf, Angerer and Theis, 2018). Cells with a high likelihood of being doublets were (doublet score >0.4) were identified with scrublet (v0.2.3, (Wolock, Lopez and Klein, 2019) and removed. Cells with <1000 UMIs or >25% mitochondrial content were removed. Genes detected (UMI>0) in less than 5 cells, marker genes, and genes on the mitochondrial genome were also removed. The final single-cell dataset contained expression measurements for 19,803 genes in 10,551 cells (donor and host), comprised of 1,488 and 1,226 cells from each MZ*nps* transplant replicate, respectively, and 7,837 cells from the WT transplanted sample.

### Data dimensionality reduction, graph construction and embedding

Count data per cell were normalized by library size (UMIs per 10,000) to account for difference in sequencing depth between cells and transformed (square-root) to account for the heteroscedasticity between genes with intrinsically different expression levels. Highly variable genes (HVGs) were identified within each sample as the top 10% of genes with the highest mean to variance ratio, and the union of HVGs in all samples defined 6,632 HVGs in the dataset. The dimensionality of HVG expression was further reduced to the top 100 PCs (explaining >99.9% of variance) which were used to calculate a cell x cell distance matrix (Euclidean) and generate a k-nearest neighbor-graph (k=5) with graphtools (v1.5.3, https://github.com/KrishnaswamyLab/graphtools). For visualization, this graph was embedded in 2-dimensions using PHATE (v1.0.11, (Moon *et al*., 2019). Embeddings were visualized using sc.pl.embedding. This graph was also used to smooth gene expression values and correct for drop out using MAGIC (v3.0.0, (van Dijk *et al*., 2018). Smoothed values were not used for differential expression testing.

### Donor cell identification

MZ*nps* knockout (KO) donor cells were distinguished from WT host cells based on the patterns of single-nucleotide-variants (SNVs) in single-cell reads per cell. In both mutant-donor samples, SNVs were called for all genes on chromosome 1 and used to define two clusters of cells using Souporcell (v2.5) (Heaton *et al*., 2020). In each sample, the cluster pertaining to MZ*nps* donor cells was identified based on enrichment of DsRed and hCD4 marker expression. As the genetic background of the WT donor and WT host cells was identical, WT donor cells were defined based on DsRed and hCD4 expression alone. Cells with >3 reads mapping to either/both DsRed and hCD4 in the WT-donor sample were annotated as donor cells. Finally, 810 KO donor cells, 3,869 WT donor cells and 5,872 WT host cells were identified.

Differentially expressed genes (DEGs) between WT and KO cells (12 hpf) were identified using the function scanpy.tl.rank_genes_groups (method=’t-test_overestim_var’, FDR < 0.05).

Differentially expressed genes between WT and KO cells at 6 hpf were identified using publicly available bulk transcriptome data deposited in the Gene Expression Omnibus (GEO) database (Riesle *et al*., 2023) (GSE162415, WT replicates n=6, KO replicates n=3). Counts per transcript were loaded into R (v4.3.2) and. aggregated per gene (sum). Genes with less than 1 count per million (cpm) in less than 2 replicates of either condition were removed, leaving 13,500 genes. Gene expression was normalized (trimmed mean of M-values, (Robinson and Oshlack, 2010), linear-models fit to each gene, and differential expression tests performed between conditions (topTreat function, LFC > 2, FDR < 0.05) using the limma R-package (v3.60.0, (Ritchie *et al*., 2015). The significance of observing the overlap between upregulated genes at 6 hpf (LFC > 2) and 12 hpf (LFC > 2) was determined using a 1-sided Fisher’s exact test. Venn diagrams were constructed using the venneuler R-package (v1.1-4). Genes were annotated as maternally deposited, downregulated in MZnps at 4 hpf or bound by NPS at 4 hpf as per (Miao *et al*., 2022). Violin plots were generated using scanpy.pl.violin.

### Identification and characterization of MZ*nps* sub-populations

After subsetting the total dataset to just the 810 KO-cells, data dimensionality reduction, graph construction and embedding was performed, as above. Leiden clustering ((Traag, Waltman and van Eck, 2019), leidenalg v0.10.0) was performed on the cell x cell graph (resolution=0.4) which identified five cell clusters. Marker genes for each sub-population were identified as those that were differentially expressed (scanpy.tl.rank_genes_groups, method=’t-test_overestim_var’, FDR < 0.05) and differentially detected (2-sided fisher exact t-test, FDR < 0.05) in each sub-population relative to cells not in the sub-population. The top markers were identified based on the combined (sum) ranking of both differential expression and detection. Significant over-representation (1-sided Fisher exact test, FDR < 0.05) of Gene Ontology biological processes (Harris *et al*., 2004), v2018) among marker genes was tested using gseapy (v1.0.6, (Fang, Liu and Peltz, 2023). Heatmaps were generated using scanpy.pl.heatmap(standard_scale=’var’).

### Developmental-age prediction and WT tissue annotation

To determine if MZnps KO cells were developmentally delayed, we embedded our dataset (query) into reference single cell dataset containing 37,183 cells collected at 12 timepoints from 3.3 to 12 hours post fertilization (hpf) (Farrell *et al*., 2018). As only cells in the 12 hpf were annotated with a tissue type in the original dataset, we first annotated all cells in the reference dataset. To annotate cells in the reference time course, we first curated a dataset spanning 3.3-24 hpf, combining cells from (Farrell *et al*., 2018; Lange *et al*., 2023; Sur *et al*., 2023), to contextualize the identification of cells with information from later developmental stages. HVGs were detected as described above and expression scaled within each dataset before datasets were concatenated. The combined expression was reduced to the top 100 PCs which were used to create a joint MNN graph of cells from all three datasets. Connections between cells from non-adjacent timepoints were masked (set to zero) in the graph kernel unless their edge weight was > 0.8. Leiden clustering (resolution=2) was performed on the masked graph and identified 37 clusters of cells. We identified marker genes between these clusters and annotated cell types based on their expression of lineage defining gene and their timepoint of collection (consistent with the known cell type present at a given developmental stage). This annotated reference was then subset to 3.3-12 hpf timepoints and used to annotate WT cells in our dataset.

HVGs in each dataset were identified as above, and HVG expression scaled within each dataset before concatenation. The dimensionality of the combined dataset was reduced to the top 37 principal components (PCs) (explaining 99.9% of the variance). Based on these PCs, the Euclidean distance between cells was calculated and used to train a k-nearest neighbor classifier (sklearn.neighbors.KNeighborsClassifier, k=5) to annotate the tissue types of WT cells in the query dataset. The distance matrix was also to create a mutual-nearest neighbor (MNN) graph between the reference and query datasets (graphtools v1.5.3) used for visualization with PHATE (v1.0.11, (Moon *et al*., 2019)).

To predict the developmental age of cells in our data, we masked query-query cell connections from the MNN-graph kernel, while preserving both query-reference and reference-reference connections. We then used MAGIC (v3.0.0, (van Dijk *et al*., 2018)) to smooth the annotated timepoint for each cell along the graph and predict the developmental age of each cell in the query and reference datasets, scaled between 3.3 and 12 hpf (the minimum and maximum timepoints in the reference dataset). Importantly, the predicted age for reference cells and WT query cells corresponded with their known timepoint of collection.

### Comparison between MZ*nps* KO sub-populations and WT tissues

The number of genes detected (UMI > 0 in at least 3 cells) was calculated for MZ*nps* KO sub-populations and WT tissues. The correlation (Pearson’s) between KO sub-populations and WT tissues was calculated based on the mean of each of the top 100 PCs within a given cell-type (scanpy.pl.correlation_matrix). Hierarchical clustering (complete) of sub-populations and tissues was performed based on 1-correlation matrix.

To determine the enrichment of WT marker genes in KO sub-populations, we first defined WT marker genes using the same marker-gene identification strategy described for the KO sub-populations above, removing genes expressed in more than 50% of cells in the total dataset. We tested for significant over representation of WT-tissue markers (1-sided Fisher’s exact test, FDR < 0.05) among the marker-genes associated for each KO sub-population. Dot plots were created in R using the ggplot2 package (v3.4.2).

Apoptotic gene expression was quantified in each cell using a gene set was curated from the GO biological processes apoptotic process (GO:0006915) and positive regulation of apoptotic process (GO:0043065). Per cell apoptosis scores were calculated by first correcting for dropout by using smoothed counts, then scaling the smoothed expression values to have zero-mean and unit variance before clipping the scaled expression at zero (to make sure genes with low expression do not have a negative contribution) and three (to ensure the score is not dominated by the high expression of a single gene). The clipped z-scores were then summed across all genes in the apoptosis gene set and the total divided by three times the number of genes in the gene set.

### Expression module (EM) analysis with GSPA

Gene-Signal Pattern Analysis (GSPA) **(Venkat *et al*., 2023)** was used to assemble modules of genes whose colocalized expression on the cell-cell graph of the dataset, which we refer to as Expression Modules (EMs). In this analysis, each gene’s expression is projected onto a library of diffusion wavelets (J=5) with varying scales and locations on the cell x cell graph (constructed above for the total dataset). The dimensionality of the resultant matrix of gene x wavelet coefficients was reduced to the top 100 PCs, a gene x gene distance matrix (Euclidean) calculated and used to create a k-nearest-neighbor gene-graph (k=5) with graphtools (v1.5.3). The distance between each gene’s wavelet coefficients and those of a constant uniform signal projected onto the diffusion wavelet dictionary was used to calculate a gene-wise localizations score - a measure of expression specificity across the cell x cell graph. Leiden clustering (leidenalg v0.10.0, resolution=2) on the gene-graph was used to identify 58 expression modules (EMs), 12 of which identified genes with a low mean localization score (<2.5) and were thus removed from downstream analyses. Module membership was calculated for each gene based on the difference in intra-module and inter-module connectivity on the gene-graph. The combined rank of each gene’s localization score and module membership used to rank each gene’s importance within its associated EM. Dot plots of top ranked genes per module were generated using scanpy.pl.dotplot. The expression of each EM in each cell was scored as described above for the apoptotic gene set. Hierarchical clustering (method=ward) based on the mean EM-expression in cells from KO sub-populations and WT tissues was performed and visualized using the python package seaborn (v0.12.2, seaborn.clustermap(standard_scale=0, row_cluster=True)).

### EM regulation in WT and MZ*nps* mutant embryos

We used publicly available datasets to examine the expression, NPS-binding profile and accessibility of promoters and enhancers of genes in each EM defined above. A publicly available timecourse measuring gene expression in WT and MZ*nps* mutant embryos between 2.5-6hpf (sampled every 30 minutes, n=2-6 per condition per timepoint, 70 samples total) was accessed through GEO (Clough and Barrett, 2016) (GSE162415). Expression per sample was library size normalized (TMM) and square-root transformed, and subset to genes with >1 count per million in at least one sample. EM expression per sample was calculated as described above with the exception that counts were not smoothed before scaling as drop out is not a significant source of noise bulk transcriptomic data. Significant differences (FDR < 0.05) in modules score between conditions at each time point were determined using scipy.stats.stats.ttest_ind. Line plots of expression were created in R using the ggplot2 package (v3.4.2). The accessibility of promoters (canonical TSS +/-500, as annotated in (Miao *et al*., 2022)) and enhancers (peak within +/-5Kb) regions for genes in each EM were analyzed using ATAC data collected from WT and and MZnps mutant embryos at 1k, dome and oblong stages of embryo development publicly available accessed through GEO (Clough and Barrett, 2016) (GSE215956, preprocessed .bw files). The mean of library depth-normalized and scaled (sklearn.preprocessing.MinMaxScaler) read counts at promoters and enhancers (within +/-200bp of summit) were calculated for genes within each EM.

NPS-binding at enhancer regions (TSS +/- 5Kb) was determined based on the intersection between the Nanog, Pou5f3 and Sox19b ChIP binding peak-regions in WT embryos as well as ATAC accessible regions in WT or mutant embryos at ∼4 hpf, as defined in (Miao *et al*., 2022). *miR-430* target sites were identified according to the presence of the seed 6-mer recognition sequence (GCACTT, (Lewis, Burge and Bartel, 2005; Giraldez *et al*., 2006) in annotated 3’ untranslated regions (3’UTRs) of genes in each EM. The statistically significant over-representation of NPS-binding and mir430 target sites was tested with gseapy (v1.0.6, 1-sided Fisher exact t-test). Heatmaps were created using the python package seaborn (v0.12.2, seaborn.heatmap).

